# Whole Body Periodic Acceleration Improves Survival and Microvascular Leak in a Murine Endotoxin Model

**DOI:** 10.1101/479477

**Authors:** Jose A. Adams, Arkady Uryash, Jose R. Lopez, Marvin A. Sackner

## Abstract

Sepsis is a life threatening condition which produces multi-organ dysfunction with profound circulatory and cellular derangements. Administration of *E.Coli* endotoxin (LPS) produces systemic inflammatory effects of sepsis including disruption of endothelial barrier, and if severe enough death. Whole body periodic acceleration (pGz) is the headward-footward motion of the body. pGz has been shown to induce pulsatile shear stress to the endothelium, thereby releasing vascular and cardio protective mediators. The purpose of this study was to determine whether or not pGz performed as a pre-treatment or post-treatment strategy improves survival in a lethal murine endotoxin model.

This study was designed as a prospective randomized controlled study in mice. pGz was performed in mice as pre-treatment (pGz-LPS, 3 days prior to LPS), post-treatment (LPS-pGz, 30 min after LPS) strategies or Control (LPS-CONT), in a lethal murine model of endotoxemia. Endotoxemia was induced with intraperitoneal injection of *E.Coli* (40mg/kg). In a separate group of mice, a nonspecific nitric oxide synthase inhibitor (L-NAME) was provided in their drinking water and pGz-LPS and LPS-pGz performed to determine the effect of nitric oxide (NO) inhibition on survival. In another subset of mice, micro vascular leakage was determined. Behavioral scoring around the clock was performed in all mice at 30 min intervals after LPS administration, until 48 hrs. survival or death. LPS induced 100% mortality in LPS-CONT animals by 30 hrs. In contrast, survival to 48 hrs. occurred in 60% of pGz-LPS and 80% of LPS-pGz. L-NAME abolished the survival effects of pGz. Microvascular leakage was markedly reduced in both pre and post pGz treated animals and was associated with increased TIE2 and p-TIE2. In a murine model of lethal endotoxemia, pGz performed as a pre or post treatment strategy significantly improved survival, and markedly reduced microvascular leakage. The effect was modulated, in part, by NO since a non-selective inhibitor of NO abolished the pGz survival effect.

## 1.0 INTRODUCTION

Sepsis is a life-threatening condition of multi-organ dysfunction caused by a dysregulated host response to infection. Septic shock is a subset of sepsis in which profound circulatory cellular and metabolic abnormalities are associated with a greater risk of mortality than sepsis alone (1). Sepsis affects more than 1.5 million humans in the USA with mortality rates of 15-30% (2). The economic burden of sepsis is highly significant. The Agency for Healthcare Research and Quality lists sepsis as the most expensive condition treated in U.S. hospitals, costing nearly $24 billion in 2013, and accounting for 6.2% of the aggregate costs for hospitalization in the USA (3). Despite hundreds of treatment trials dating to the 1960s, interventions to improve survival from sepsis have not significantly lowered mortality.

Administration of lipopolysaccharide (LPS), the endotoxin derived from the purified outer membrane of *E. Coli*, produces systemic inflammatory effects of sepsis in mice. Exposure to LPS causes a dose-dependent activation of a widespread cascade of inflammatory mediators that disrupt the endothelial barrier. This leads to intracellular hyperpermeability, multiple organ dysfunction and if sufficiently severe, death (4). Such a situation calls for an effective, prompt, endothelial repair strategy that has not yet been promulgated. Large quantities of nitric oxide that are released into the circulation through the action of inducible nitric oxide synthase (iNOS) are an important component of this inflammatory cascade in sepsis disruption to the endothelial barrier. In contrast, small quantities of nitric oxide normally released from endothelial nitric oxide (eNOS) by flowing blood are a crucial determinate of inter-endothelial junctions (5). The potential effectiveness of eNOS was reported by Yamashita et al almost 20 years ago in which chronic overexpression of endothelial derived NO by transgenic mice resulted in resistance to LPS-induced hypotension, lung injury and death (6).

In this paper, we employed non-invasive, periodic acceleration (pGz), a means to increase pulsatile shear stress to the endothelium a phenomenon that also takes place during exercise to stimulate increase release of NO into the circulation as an alternative to overexpression by transgenic nice. This was accomplished by rapidly and repetitively moving a mouse in headward-footward direction to induce pulsatile shear stress to the endothelium (7–9). We and others reported that pulsatile shear increases expression of both endothelial derived nitric oxide synthase (eNOS) and neuronal derived nitric oxide synthase (nNOS) both which are produced in nanomolar concentrations, and are important in modulating the anti-inflammatory response in sepsis (9–11). We have previously shown in animal models of whole body and focal ischemia reperfusion injury of the heart, brain, and skeletal muscles, that pGz improves outcomes, (12–19). In part, the effects of pGz are related to increased release of eNOS into the circulation as well as prostaglandins, adrenomedullin, and signaling via Phosphoinositide 3-kinase protein kinase B pathway (PI3K-AKT). pGz also reduces intracellular calcium overload at the cellular level. (10, 11, 19-22).

We hypothesized that pGz performed as a pre-treatment or post-treatment strategy in a lethal murine endotoxin model might confer improved survival.

## 2.0 MATERIALS AND METHODS

### 2.1 Animal Preparation & pGz

This study protocol was approved by the Institutional Animal Care and Use Committee of Mount Sinai Medical Center and conforms to the Guide for the Care and Use of Laboratory Animals published by the National Institutes of Health (NIH Publication No. 85-23, revised 1996) Protocol No. 17-20-A-04.

The motion platform that imparts pGz has been previously described (18, 23). The platform moves the horizontally placed body in the z-plane. The frequency of periodic acceleration has been previously determined in our laboratory for mice to be 480cpm. The mice were acclimated to a mouse holder (Kent Scientific Design, Torrington, CT) for 2 days and thereafter the mice voluntarily walked into the mice holder. The mice holder was placed on the pGz motion platform. pGz was carried out at a frequency of 480 cycles/minute and acceleration in the z-plane (Gz) of ± 3.9 m/sec^2^

### 2.2 Experimental Design

Survival Experiments: Prior to LPS inoculation, mice were randomly assigned to one of three groups; a) Pre-treatment (pGz-LPS) (n=12) pGz was performed 1 hrs. per day for 3 days, b) Post-treatment (LPS-pGz) (n=12) pGz was performed starting 30 min after LPS and continued for 1 hrs., c) LPS Control (LPS-CONT) (n=12). Mice received an intraperitoneal injection of *E.Coli* lipopolysaccharide (Sigma Aldrich, St. Luis, MO) at a lethal dose of 40mg/kg diluted in phosphate buffered saline, total volume 0.1ml. A separate group of mice received same volume of phosphate buffered saline but did not receive LPS or pGz and were used as Sham (n=8) controls.

Behavioral scoring was performed in all animals for the initial 48 hrs. after LPS injection. The Behavioral Scoring criteria utilized has been described by Shrum et al (24).

### 2.3 Nitric Oxide Inhibition

A separate group of animals received a non-specific nitric oxide synthase inhibitor (L-NAME, 1.5mg/ml) in their drinking water for 5 days and were randomized prior to LPS injection (same dose as above) to receive pGz pre-treatment (n=8) post-treatment(n=8) or LNAME-Control (n= 8).

### 2.4 Determination of Microvascular Leakage

LPS increases microvascular permeability, with endothelial tight junction disruption, and increased endothelium permeability. To determine the severity of microvascular leakage and the effects of pGz on such, a separate group of animals was injected intravenously with 0.5% sterile filtered Evans Blue at a dose of 8ml/kg (total volume 0.2ml) 5 hrs. after LPS. The animals were randomized to LPS-CONT (n=8) pGz-LPS (n=8), LPS-pGz (n=8), or Sham (received Evans Blue but did not received LPS or pGz, n= 4). At 6 hrs., the animals were sacrificed and Evans Blue extravasation was quantitated using SpectraMax Plate Reader (Molecular Devices, Sunnyvale, CA) (25). Microvascular leakage was determined in the lungs, liver and mesenteric circulation (testis).

### 2.6 Protein Expression

Protein extraction was performed as previously described. Protein analysis was performed using Western Blot method and visualized by enhanced chemifluorescence. (S1 Supplemental Information)

### 2.7 Euthanasia

After completion of each of the experimental protocols animals were euthanized by a method approved by the American Veterinary Medical Association Guidelines on Euthanasia. (S1 Supplemental Information)

### 2.8 Statistical Analysis

All values are reported as means ± SEM. Continuous variables were evaluated by analysis of variance for repeated measures. For variables with significant differences, post hoc analysis was done using Tukey HSD for equal or unequal sample size. Comparisons of discrete variables were evaluated by Fisher’s exact test. Statistical analyses were performed using STATISTICA (StatSoft Inc., Tulsa, OK). Sample size was calculated using Statistica based on power analysis with α=0.05 and power 0.80. A p value of < 0.05 was considered statistically significant.

## 3.0 RESULTS

### 3.1 Behavioral Scores & Survival

Using the behavioral scoring criteria of Shrum et al (24), there were significant differences in outcomes between LPS-CONT compared to pre-treated (pGz-LPS) and post-treated (LPS-pGz) 8 hrs. after LPS (Fig 1). At 30 hrs. after LPS injection, nontreated animal had 100% mortality. In contrast, pGz-LPS and LPS-pGz had 60 and 80% survival (p < 0.001). The survival benefit of pGz persisted beyond 48 hrs. without additional pGz treatment. L-NAME (non-selective NO inhibitor) significantly reduced survival for both non treated and pGz treated animals. Survival for LNAME animals did not surpass 16 hrs. (Fig 1).

**Fig. 1:**
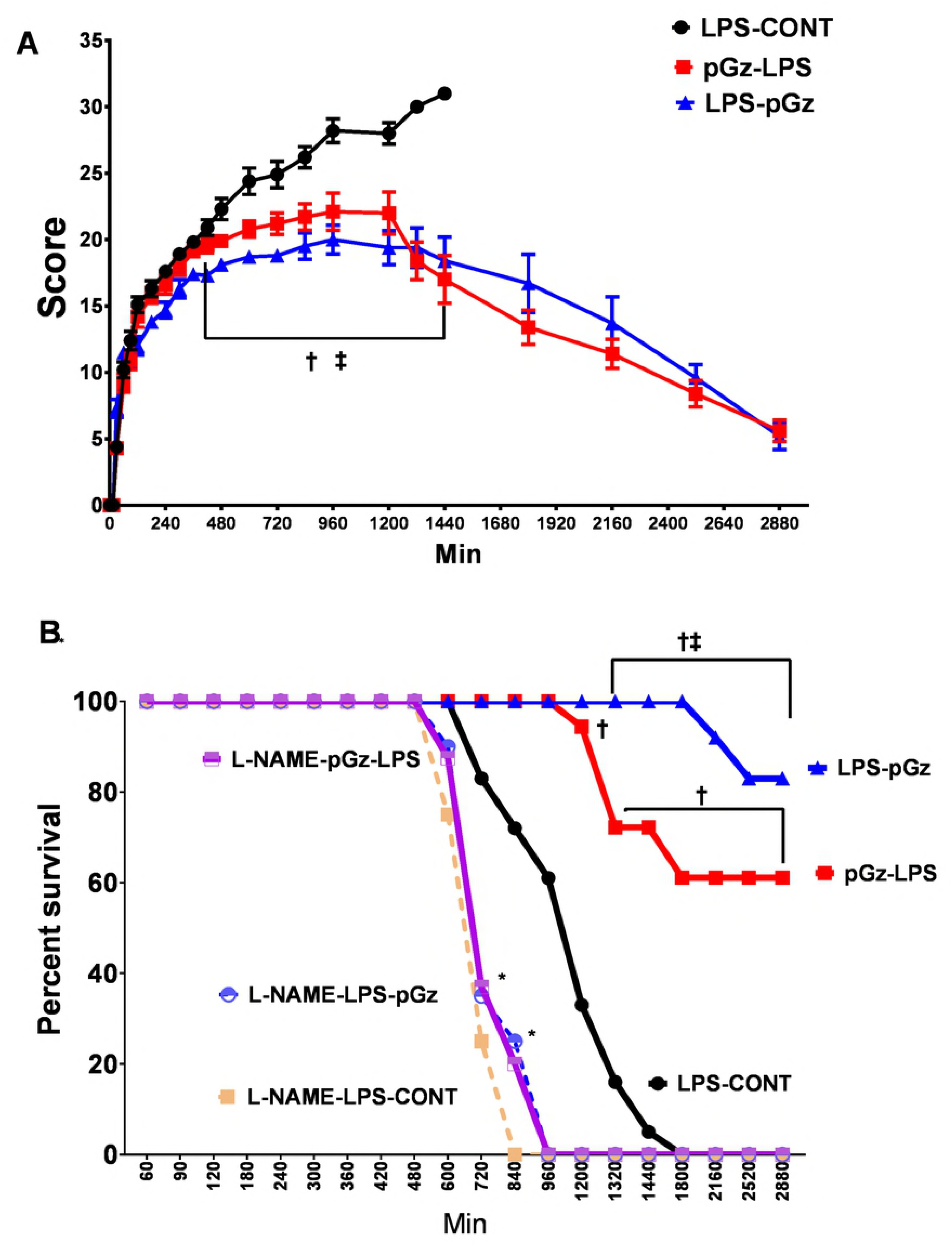
Behavioral Scoring and Survival Kaplan-Meyers Curve in Mice After Lethal Dose of LPS. **(A)** Mean behavioral scores in mice every 30 mins, using scoring criteria of Schrum et al (24) in LPS-CONT, pGz pre-treatment (pGz-LPS) and pGz post-treatment (LPS-pGz). Scoring was begun 30 min after LPS injection and continued until death or survival for up to 48 hrs. **(B)** Survival in mice exposed to a lethal dose of LPS over the initial 48 hrs. Fifty percent survival in LPS treated mice was reached at 20 hrs. after LPS in control animals (LPS-CONT), in contrast all LNAME pretreated animals reached 50% survival at 12 hrs.. 80% of LPS-pGz and 60% of pGz-LPS animals survived beyond 48 hrs.. Data are mean ± SEM (**†**p<0.01 pGz-LPS or LPs-pGz vs. LPS-CONT **ǂ** p< 0.01 LPS-pGz vs. pGz-LPS, * p< 0.01 L-NAME-LPS-pGz or L-NAME-pGz-LPS or L-NAME-LPS-CONT vs LPS-CONT)

### 3.2 Microvascular Leakage and TIE2 Expression

LPS induced a significant increase of at least double the amount of microvascular leakage determined in Sham animals (which received Evans Blue, but did not receive LPS or pGz) in mesenteric, lung and liver vasculature. pGz pre and post treatments reduced microvascular leakage in all three vascular beds by 50% (Fig 2).

**Fig. 2:**
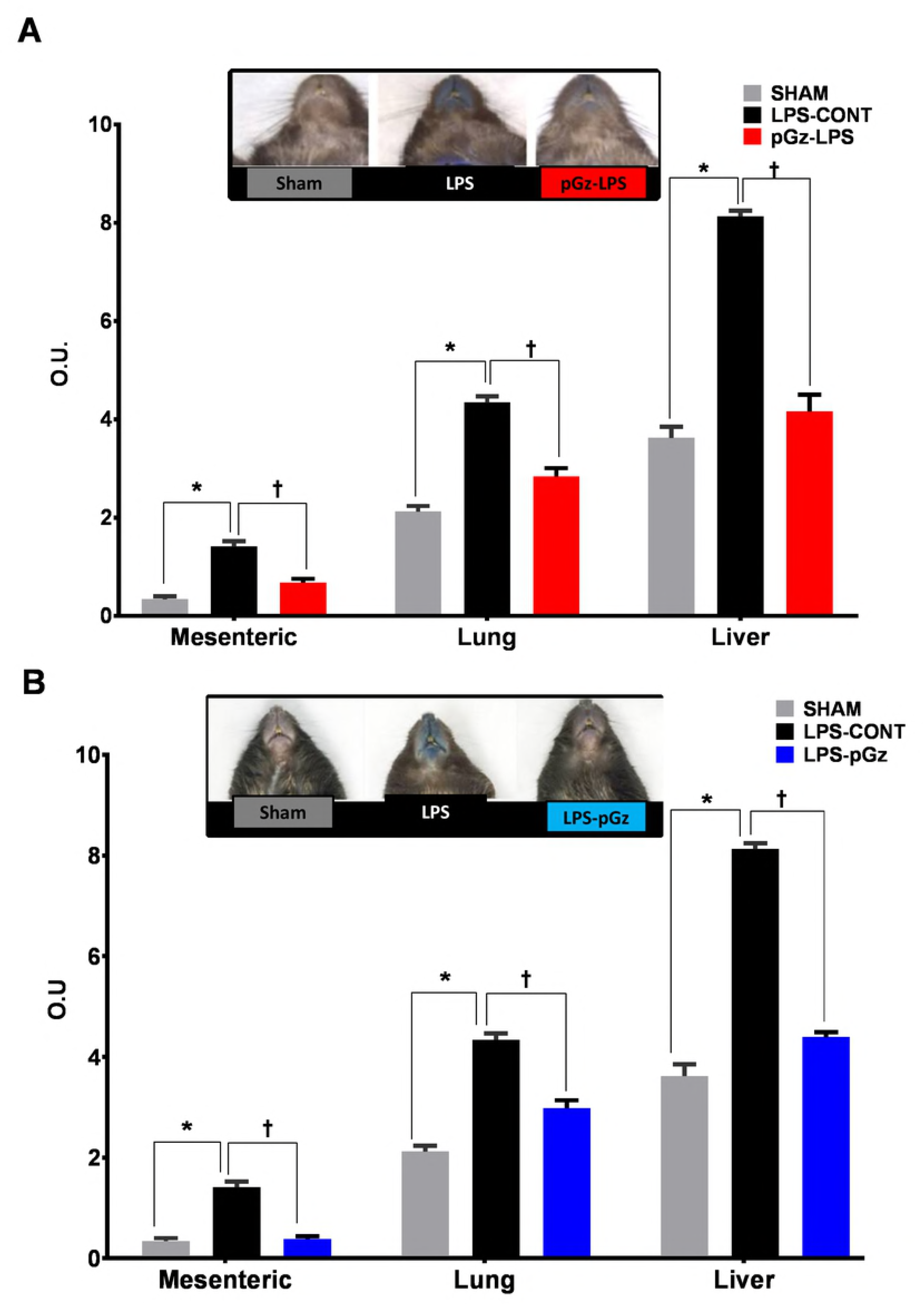
Microvascular Leakage after LPS in Mesenteric, Lung and Liver Vasculature. Mesenteric, lung and liver microvascular leakage determined by spectrophotometric optical density (OD) of Evans Blue, in Sham, LPS-CONT and pGz pretreated (pGz-LPS) **(A)** and **(B)** post-treated mice (LPS-pGz) mice. LPS-CONT (n=8) pGz-LPS (n=8), LPS-pGz (n=8), or Sham (received Evans Blue but did not received LPS or pGz, n= 4). Data are mean ± SEM. (* p<0.01 vs. Sham, Ɨ p<0.01 vs. LPS-CONT)

Expression of TIE2 was unchanged with LPS but both pre and post treatment with pGz increased TIE2 by 150%. In contrast, p-TIE2 decreased after LPS by 50% and pre and post treatment with pGz increased the latter to Sham levels (Fig 3).

**Fig. 3:**
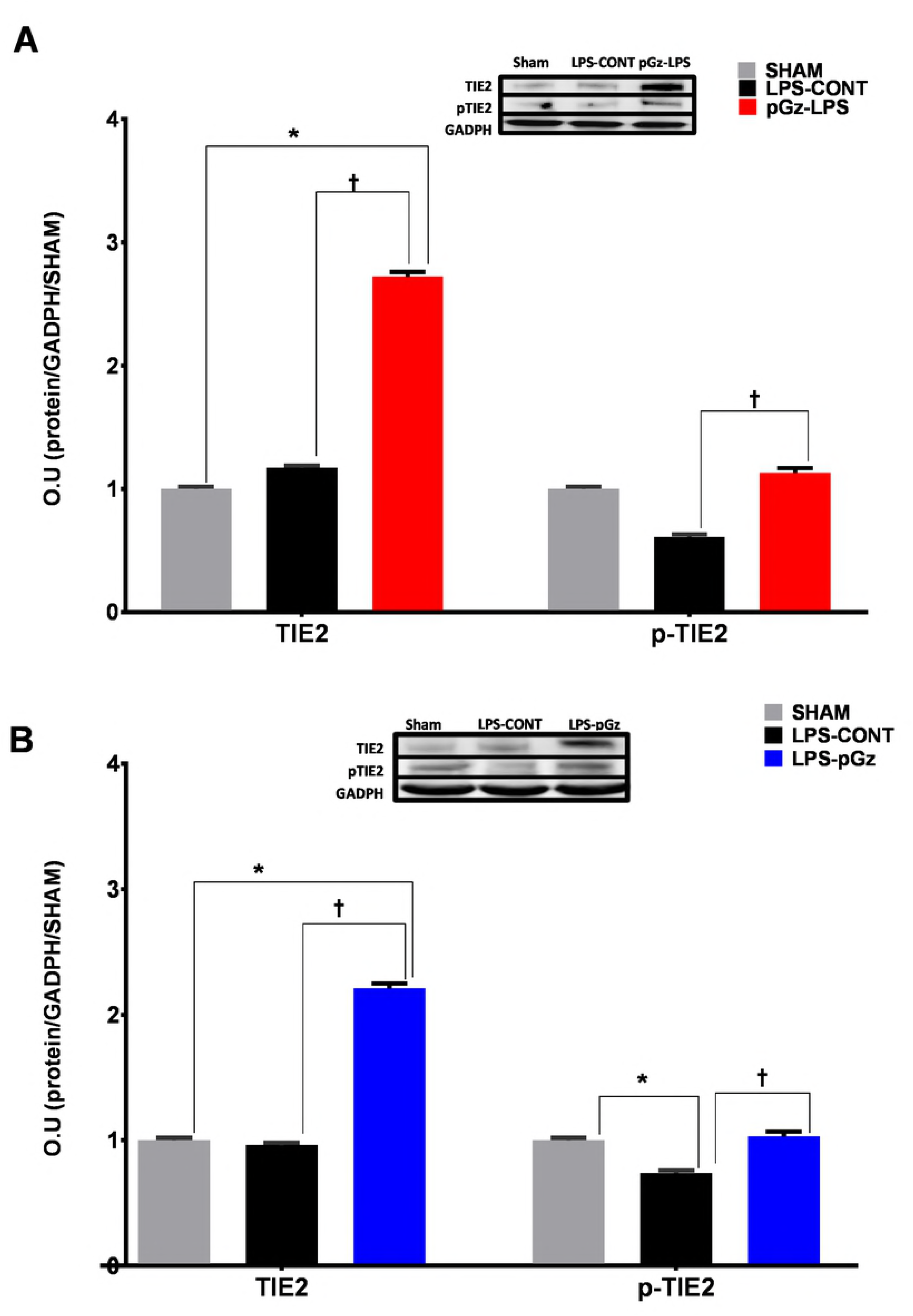
Expression and phosphorylation of TIE2 in Mice after LPS. Expression of the tyrosine kinase receptor TIE2 and its phosphorylation in mice, 6 hrs. after LPS. LPS did not significantly change expression of TIE2, but significantly decreased TIE2 phosphorylation. **(A)** pGz pre-treatment (pGz-LPS) and **(B)** pGz post-treatment (LPS-pGz) significantly increased both TIE2 expression and phosphorylation. Inserts are representative western blots of protein expression of TIE2, p-TIE and GADPH in Sham, LPS-Control, and pGz-LPS and LPS-pGz. LPS-CONT (n=8) pGz-LPS (n=8), LPS-pGz (n=8), or Sham (received equal volume of phosphate buffer but did not received LPS or pGz, n= 4). Data are mean ± SEM (* p<0.01 vs Sham, Ɨ p<0.01 vs. LPS-CONT).

## 4.0 DISCUSSION

The present study demonstrates that pGz as a pre or post treatment strategy significantly improves survival in a lethal model of LPS induced endotoxin shock. To our knowledge, our study is the first to show that repetitive passive motion of the body as produced by pGz, markedly improves endotoxin shock survival accompanied by a decrease of microvascular leakage in lungs, liver and mesenteric vasculature.

Interventions such as exercise and remote ischemic pre and post conditioning have been used as therapeutic modalities in mice models of LPS induced inflammation. Table 1 summarizes exercise or remote ischemic pre or post treatment interventions with doses of LPS and survival outcomes in all studies which have reported at least 24 hrs. survival in mice.

**Table 1:** Comparison of studies using non pharmacological pre or post treatment strategies in LPS induced sepsis in mice

| Author | Year | Species | Dose of LPS | Pre or Post Treatment | Mode | 24 hr Survival | 48 hr Survival |
| --- | --- | --- | --- | --- | --- | --- | --- |
| Ishizashi et al(28) | 1995 | 5 week old C57BL/6J | IV LPS 20mg/kg | Pre-Treated (P. acnes, IP 7 days before exercise). Exercise 120 min prior to LPS | Voluntary Wheel Running | Control =0<br>Exerc= 22% |  |
| Martin et al (27) | 2013 | 22 month C57BL/6J | Max Dose IP 0.33mg/kg | Pre treatment 10 weeks | Voluntary wheel running | Control=73%<br>Exerc=100% |  |
| Kim et al (29) | 2014 | 6 weeks mice BALB/c | IP 20 mg/kg | Remote Ischemic Pre (RIPC) & Post Conditioning (RpostC), immediately before or after LPS | Hind-limb tourniquet 3 cycles (10min Ischemia/10min reperfusion) | Control=80%<br>RIPC =100%<br>RpostC =90% | Control=15%<br>RIPC =65%<br>Rpostc =65% |
| Irahara et al (26) | 2016 | 11 weeks C57BL/6J | Max Dose IP 10mg/kg | Post Treatment exercise | Treadmill 30 min X 3 after LPS and 30 min X 2 next day | Control=80%<br>Exercise=100% | Control=60%<br>Exercise=100% |
| Adams et al<br>Current work | 2018 | 6-10 weeks C57BL/6J | IP 40mg/kg | Pre and Post Treatment | pGz<br>Pre Treatment =3 days<br>Post Treatment =30 min after LPS | Control= 6%<br>pGz-LPS= 75%<br>LPS-pGz= 100% | LPS-Control= 0%<br>pGz-LPS=61%<br>LPS-pGz=83% |

Compendium of mice studies using non pharmacological interventions pre or post treatment in models of LPS induced endotoxin in mice published from 1995 to 2018 with at least 24 hrs. survival. Each study shows the dose of LPS, method of administration (IP=intraperitoneal, IV=intravenous) the pre or post treatment strategy, the modality and 24 and 48 hrs. survival.

Only one study utilized exercise as a post-treatment strategy, viz., treadmill exercise during the acute phase after LPS up to 72 hrs. (26). Survival in sedentary controls at the highest dose of LPS (10mg/kg) was 50%, whereas 100% survived in low dose LPS control and exercise groups. The highest dose used by this investigation was 25% of the LPS dose used in our study. Based on our behavioral score data it is obvious that exercise after a lethal dose of LPS such as the one used in our experiments cannot be performed because of the severity of symptoms. Our data is consistent with other investigators who have also shown improved survival when animals are pre-treated with an active exercise intervention (27–29).

Survival was significantly decreased in L-NAME treated animals. The latter was not surprising since other animal studies have also shown similar findings with non-selective NOS inhibition (30, 31). Human studies performed with nonselective NOS inhibition also support the latter (32, 33). We have previously shown the time course of eNOS upregulation and phosphorylation in both animal models and cells under pGz (11, 23). Data are conflicting with regards to eNOS and survival and outcomes from LPS inflammatory response and sepsis induced by cecal ligation. Some studies using eNOS deficient mice demonstrate that eNOS deficiency decreases mortality and improves outcomes from LPS sepsis (34, 35) while others suggest the opposite (6, 36-38). All three NOS isoforms play a role in sepsis, however, the extent timing, tissue location and eNOS uncoupling are all important to take into consideration (6, 34-37, 39-41). eNOS has been shown to modulate inflammatory response, and particularly that of NFĸβ and its crosstalk with iNOS (42). The effects of eNOS on other cytokines also remains to be determined.

It is important to acknowledge that a single plausible mechanism for our findings is not possible, since pGz produces a multifaceted response of cytoprotective proteins, including antioxidant defense (23, 43).

Microvascular leakage has been described in various models of sepsis including LPS and its etiology multifactorial, with endothelial dysfunction and disruption of tight junctions via various mechanisms (44, 45). We have shown that pGz both pre and post treatment decreases microvascular leakage induced by LPS. The decrease in microvascular leakage by pGz is also likely multifactorial. Microvascular leakage has been shown by others to be decreased by eNOS. (46). We also found that pGz produces a significant increase in Tie2 and restoration of p-Tie2. Tie2 is a tyrosine receptor kinase which has been shown to be important in maintenance of tight junctions and decreasing vascular hyperpermeability and specifically during sepsis (47, 48). Tie2 stimulation can promote a broad range of microvascular anti-leakage including endothelial activation. David et al showed a decrease in total Tie2 expression and phosphorylation in LPS treated mice and a restoration and improved survival in mice pretreated with a known Tie2 agonist (vaculotide) (49). The latter has been shown to improve endothelial tight junctions. Oxidative stress has been shown by others to be a hallmark of the inflammatory response, and to play a role in microvascular leakage. We did not specifically studied ROS production in this study but we have previously shown that pGz in mice also increases antioxidant enzymes (SOD, Catalase) and total antioxidant capacity (23).

### Potential Clinical Application

Increase pulsatile shear stress to the vascular endothelium by way of mechanical addition of pulsations is a novel method with clinical applicability. A motion platform to produce Whole Body Periodic Acceleration in humans was formerly available but is no longer being marketed (NIMS, Miami FL 33137) However, it was not portable because it weighed 179 kg and was difficult to perform nursing care while subject was moving back and forth.

A new non-invasive, portable, passive simulated jogging device (JD) has recently been fabricated and tested in normal and hypertensive humans as a clinical trial with favorable benefits. (50, 51) This device utilizes microprocessor controlled, DC motorized movements of foot pedals placed within a chassis to repetitively tap against a semi-rigid surface for simulation of locomotion activities while the subject is seated or lying in a bed (Gentle Jogger®, Sackner Wellness Products LLC, Miami FL 33132). It weighs about 4.5 kg with chassis dimensions of 34 × 35 × 10 cm. It is placed on the floor for seated applications and secured to the footplate of a bed for supine applications. Its foot pedals rapidly and repetitively alternate between right and left pedal movements to actively lift the forefeet upward about 2.5 cm followed by active downward tapping against a semi-rigid bumper placed within the chassis. In this manner, it simulates feet impacting against the ground during selective speeds of locomotor activities. Each time the moving foot pedals strike the bumper, a small pulse is added to the circulation as a function of pedal speed that produce up to 190 steps per minute, thereby increasing pulsatile shear stress to the endothelium with its benefits as in the current mice study.

The maximum acceleration forces in seated posture from a triaxial accelerometer during run speed operation of JD was ±5.4 m/s^2^ over tibia, ±5.1 m/s^2^ over femur, and ± 1.0 m/s^2^ over forehead. The maximum acceleration forces in supine posture from a triaxial accelerometer during run speed operation of JD was ±2.9 m/s^2^ over tibia, ±6.3 m/s^2^ over femur and ±3.6 m/s^2^ over the forehead. Clinical trials for prevention and treatment of human sepsis will be needed to demonstrate its effectiveness of this safe low risk modality according to FDA wellness guidelines.

### Study Limitations

We must also acknowledge what others have previously recognized and thoroughly reviewed about the LPS murine model and its differences between it and other murine models of sepsis and clinical sepsis (52–54). This study addresses short term survival and microvascular leakage, in LPS induced septic shock and long term outcomes could not be addressed The LPS dose utilized in our experiments was specifically selected due to the severity of injury produced, thereby providing a model with very high mortality. We utilized and orally administered dose of L-NAME, a nonspecific NO inhibitor, since specific eNOS inhibitors in mice are lacking. It can be argued that the exact dose of L-NAME each animal received may be slightly different, however the effects of NO inhibition on survival from LPS were uniformly dismal. We also did not compare pGz to any other intervention, since there are no other passive exercise strategies suitable for comparison in mice with this level of LPS induced injury.

Notwithstanding the above limitations, we conclude that pGz is a simple method which increases survival, and reduces microvascular leakage in a mouse model of LPS induced endotoxin shock. pGz may serve as an adjunctive therapeutic strategy to current pharmacological modalities to ameliorate or improve outcomes from sepsis. Human clinical trials are needed to confirm efficacy in human sepsis.

## ACKNOWLEDGMENTS

This study was supported in part by a grant from the Florida Heart Research Institute

